# The pyriproxyfen metabolite 4’OH- pyriproxyfen disrupts thyroid hormone signaling and enhances Musashi-1 levels in neuroprogenitors

**DOI:** 10.1101/352088

**Authors:** Petra Spirhanzlova, Sébastien Le Mével, Karn Wejaphikul, Bilal Mughal, Jean-David Gothié, Anthony Sébillot, Lucille Butruille, Michelle Leemans, Theo Visser, Sylvie Remaud, Jean-Baptiste Fini, Barbara Demeneix

## Abstract

Epidemiological and experimental studies have raised questions as to whether the insecticide pyriproxyfen (PPF) could be implicated in the increased incidence of microcephaly associated with ZIKA infection during pregnancy. This pesticide is documented as a thyroid hormone (TH) disrupting chemical. We investigated whether environmentally relevant amounts of its main metabolite, 4’-OH-pyriproxyfen (4’-OH-PPF), modified TH signaling and early neuronal development. First, an *in silico* study revealed strong affinity of 4’-OH-PPF to fit the ligand binding pocket of TH receptors (TRs). Further, *in vitro* assays on human cell lines showed 4’OH-PPF (> 3 mg/L) to act as a TRα antagonist. Next, using a transgenic Xenopus TH-sensitive reporter system, Tg(*thibz*:GFP) tadpoles showed that 4’OH-PPF (> 10^−7^ mg/L) displayed TH-disruptive activity and reduced tadpole mobility (> 10^−1^ mg/L). Exposure to 4’OH-PPF significantly reduced Xenopus head size at levels equivalent to the maximum recommended daily intake of PPF (3× 10^−1^ mg/L). Most strikingly, in both the Xenopus system *in vivo* and in mouse neurosphere cultures, environmentally relevant concentrations of 4’OH-PPF increased expression of the gene encoding an RNA-binding protein that enables ZIKA replication: Musashi-1 (*msi1*) in neurogenic brain areas. We conclude that first, the PPF metabolite, 4’OH-PPF, disrupts thyroid signaling, neuronal development and behavior in Xenopus embryos, and second, that it increases Musashi-1 levels in neurogenic zones of both mouse and Xenopus, creating the potential to enhance viral replication. As PPF is used in areas with high microcephaly incidence and is readily broken down to 4’OH-PPF, these findings provide a plausible mechanism whereby PPF could, through modulating expression of Musashi-1, exacerbate the effects of ZIKA virus infection.

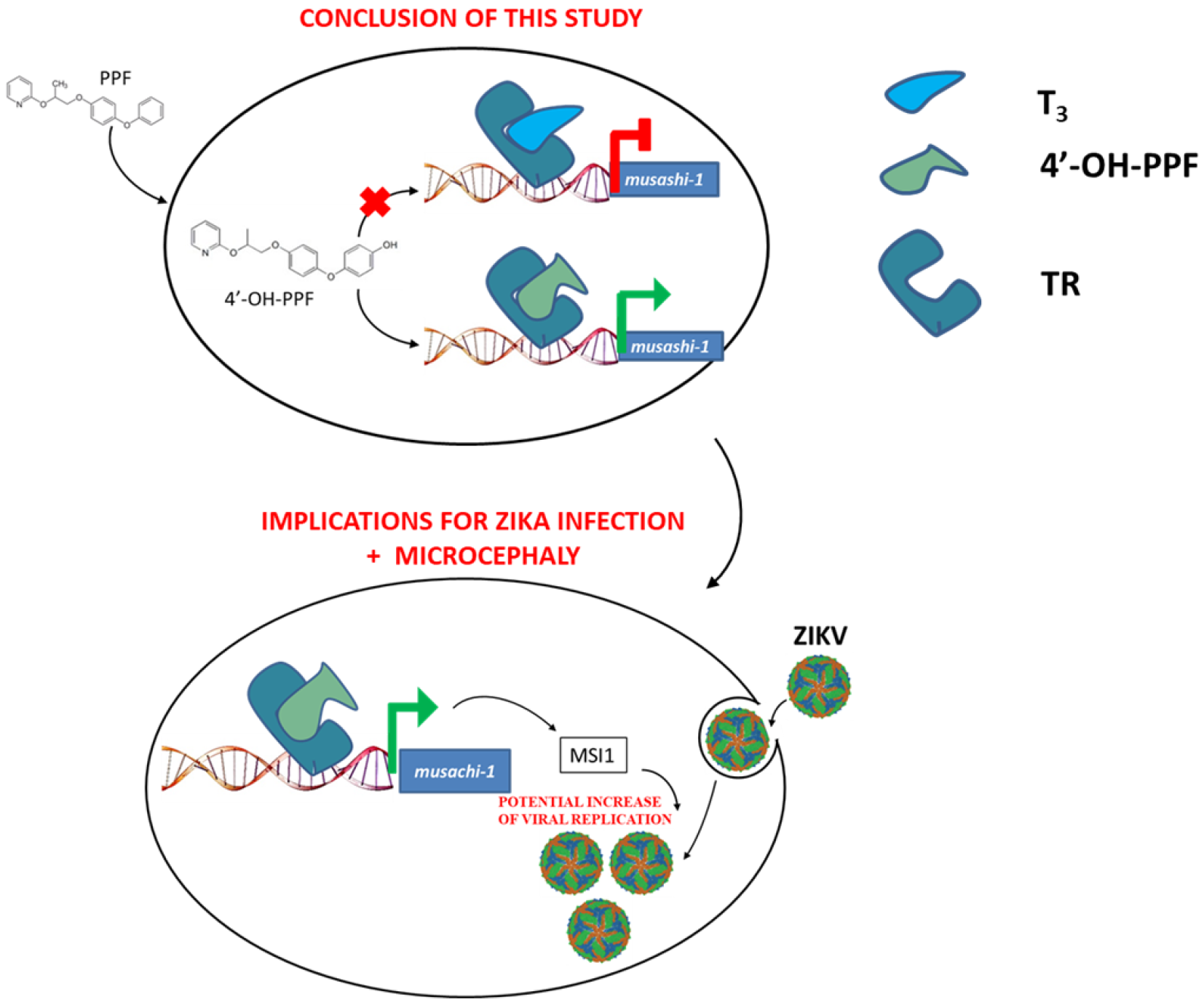

## Introduction

A number of lines of evidence link infection by the ZIKA virus to the increased incidence of microcephaly in Central and South America starting from early 2016 (Alvarado and Schwartz, 2017; de Araújo et al., 2016; Faizan et al., 2016; de Oliveira et al., 2017). However, many authors have queried as to whether other environmental factors could contribute to the increased incidence of microcephaly (Butler, 2016a, 2016b; Rodrigues and Paixao, 2017). Certain authors have queried whether the use of the pesticide pyriproxyfen (PPF) could be implicated (de Albuquerque et al., 2016; Evans et al., 2016; Parens et al., 2017; REDUAS, 2016). PPF was first introduced into Brazilian drinking water in late 2014 as a mean of controlling the *Aedes aegypti* mosquito population, the vector for the ZIKA virus. Highest amounts of PPF were used in the north-east, but whether or not the pesticide had any toxic effects, related or not to microcephaly was not thoroughly investigated, neither epidemiologically nor experimentally until recently (Parens et al., 2017).

PPF is a juvenile hormone analog primarily used against household insects including fleas and on agricultural crops and its use has been approved worldwide (US EPA, 2011; WHO, 2007; Wilson, 2004). WHO recommends that daily intake of PPF should not exceed 0.3 mg/L for an average adult and the recommended concentration of PPF in drinking water containers is 0.01 mg/L (WHO, 2007). The use of PPF is not limited only to South America. PPF has been added in the drinking water in Cambodia or Malaysia and it has been found in Spanish river water and fish at concentrations reaching 90 ng/L and 0.1-0.3 ng/g respectively (Belenguer et al., 2014; Invest and Lucas, 2008). However, PPF has been documented as interfering with thyroid hormone (TH) signaling (Wegner et al., 2016) an essential hormone for brain development (Bernal, 2005).

The major pathway of PPF metabolism is hydroxylation at the 4’-position, producing majoritarily 4’-OH-PPF in Xenopus, mice, rats, goats, house flies and chicken (Fujimori, 1999; Ose et al., 2017; Yoshino et al., 1995; Zhang et al., 1998). To our knowledge, no studies have yet assessed the toxicological potential of 4’-OH-PPF.

Given the lack of information on 4’-OH-PPF, we used *Xenopus laevis* tadpoles, a well-known model for testing TH disruption (Fini et al., 2012, 2017) and mouse neurospheres to investigate the potential of PPF and 4’-OH-PPF to disrupt thyroid hormone (TH), signaling during brain development. We also examined how PPF and 4’-OH-PPF affected expression of the key protein, Musashi-1 known to be required for ZIKA replication (Chavali et al., 2017). *Musashi-1* is down regulated by TH in the neurogenic zone of the adult mouse brain (López-Juárez et al., 2012), one example of the multiple processes implicating TH during the development of the central and peripheral nervous system, from stem cell proliferation, lineage decisions, differentiation, migration, synaptogenesis and myelination (Bernal, 2005; Morreale de Escobar et al., 2004). Impaired thyroid signaling during development results in brain defects in human (Korevaar et al., 2016; Pop et al., 1999).

## Materials and methods

### 1. Materials

#### 1.1 Reagents

3,3′,5-Triiodo-L-thyronine sodium salt (T_3_) (Sigma - CAS 55-06-1), 4’OH pyriproxyfen (HPC – Ref. 675366), Pyriproxyfen (Sigma – CAS 95737-68-1), Dimethyl sulfoxide –DMSO (Sigma - CAS 67-68-5), DMEM/F12 (Gibco), Ethyl 3-aminobenzoate methanesulfonate salt (MS 222) (Sigma - CAS 886-86-2), NH3 (synthetized by AGV Discovery-France), Sodium bicarbonate (Sigma-CAS 144-55-8), Paraformaldehyde (Sigma – CAS 30525-89-4), Succrose (Sigma- CAS 57-50-1), Ethylen glycol (Sigma- CAS 107-21-1), Tris, pH 8.0, KCl, MgCl_2_, glycerol, DTT (Sigma), [^125^I]T_3_, Agilent RNA 6000 Pico Kit, Reverse Transcription Master Mix (Fluidigm), Power SYBR Master Mix (Life technologies - ref.4368708), RNAqueous–micro kit (Life technologies - ref. AM1931), Primers (Eurofins), TNT^®^ T7 Quick Coupled Transcription/Translation System (Promega, Leiden, The Netherlands), nitrocellulose membrane (Millipore HA filters, 0.45 μm), BSA, Normal goat serum, Papain, Cysteine and DNAse, FGF2 (Peprotech), TaqMan Preamp MasterMix, TaqMan Universal PCR MasterMix (both ThermoFisher), specific TaqMan probes (*Msi1*, Mm00485224_m1; *ActB*, Mm01205647_g1; *Gapdh*, Mm99999915_g1; *Hprt*, Mm00446968_m1), Msi1 primary antibody (Rabbit polyclonal to Msi1- Abcam, UK), Alexa Fluor 488-conjugated anti-rabbit antibody (Thermofisher Scientific), Milli-Q reference water, Nuclease free water, Evian water in glass bottle 75 cL or another mineral water with reproducible quality, Liquid nitrogen.

#### 1.2 Equipment

Electrical devices: BioAnalyzer (Agilent), Daniovision (Noldus), Fluorescent microscope equipped with 25× objective and long pass GFP filter – Olympus, Incubator, NanoDrop ThermoScientific, QuantStudio 6 flex QPCR machine (Life technologies), PCR machine (Biorad), Vortex mixer.

Laboratory utensils: 384-well hardshell plate clear, 50 mL Greiner capped tubes (Greiner Bio-one - ref.227261), 96-well black, conic well plate (Greiner Bio-one - ref. 651209), Adhesive covers for 384 well plates, Sterile dissecting tools, Microcentrifuge tubes 0.2-1.7 mL, flip top, Multi-distribution pipette, PCR tube Strips 0.2 mL Eppendorf, Micropipettes, Pipette tips 0.2-5000 μL, Transfer pipets with extended fine tip, Transparent flat 6-well plates (TPP, Switzerland), Transparent flat 12-well plates (TPP, Switzerland).

#### 1.3 Software

Capture pro (QImaging), ImageJ – optional, GraphPad Prism 7, MS Excel, QuantStudio™ Real-Time PCR Software, Ethovision XT 11.5, Qiagen CLC Drug discovery 3, NHR scan.

### 2. Methods

#### 2.1 In silico

Qiagen CLC Drug discovery 3 was used to assess ligand-protein interactions. Human crystal structure of thyroid hormone receptor alpha (TRα, 4LNW), bound to a ligand was used. The list of compounds including pesticides and insecticides used have been outlined in supplementary table 1 with their corresponding PubChem compound id. Thyroid hormone antagonist NH3 (Lim et al., 2002) and 4’-OH-PPF were drawn manually. 10,000 iterations were run for each ligand-protein interaction, against the T_3_ binding pocket of TRα, and top 10 recurring interactions were analysed. The “score” represents the steric fit of ligand in the T_3_ binding pocket, where a more negative score indicates a higher ligand-protein interaction. Mean of top 10 interactions scored were ranked and displayed in supplementary table 1.

#### 2.2 Cell culture, transfection, and luciferase assays

JEG3-cell culture and transfection were performed as previously described (van Gucht et al., 2016). Briefly, FLAG-tagged human TRα1 was overexpressed in JEG3 cells together with a DR4-TRE luciferase reporter construct and pMaxGFP as a transfection control. After 24 hours, cells were incubated for 24 hours in DMEM/F12+0.1% BSA containing 1 nM of T3 and the indicated concentrations of 4’-OH-PPF or the TRα1-antagonist, NH3. Luciferase activity was measured in cell lysates as previously described (van Mullem et al., 2012). The ratio of the luciferase and GFP activities was calculated to correct for transfection efficiency. The results are expressed as percentage of the ratio at 0 nM 4’-OH-PPF or NH3. Data from two independent experiments performed in triplicate are shown as mean±SEM and analysed by One-way ANOVA with Tukey’s post-test. Statistical significance was considered when p-values < 0.05.

#### 2.3 [^125^I]T3 competitive binding assays

FLAG-tagged TRα1 protein was synthesized by the TNT^®^ T7 Quick Coupled Transcription/Translation System (Promega, Leiden, The Netherlands), according to the manufacturers’ protocol. 0.25 μL of TRα1 protein was incubated for 2 hours at 30°C in binding buffer (20 mM Tris, pH 8.0, 50 mM KCl, 1 mM MgCl_2_, 10% glycerol, 5 mM DTT) containing 0.02 nM of [^125^I]T_3_ and 0-10,000 nM of the unlabelled competitive compound (T_3_, 4’-OH-PPF, or NH3). Subsequently, samples were filtered through a nitrocellulose membrane (Millipore HA filters, 0.45 μm). The amount of TR-bound [^125^I]T_3_ on the membranes was quantified and expressed as percentage maximal [^125^I]T_3_ binding for each competitor. The [^125^I]T_3_ dissociation curve and the dissociation constant (Kd) were computed using GraphPad Prism version 7.0 (GraphPad, La Jolla, CA). The data were shown as mean±SEM of at least two independent experiments performed in duplicate.

#### 2.4 Primary mouse NSC neurosphere cultures

C57BL/6 wild-type male mice, eight weeks old, were purchased from Janvier (Le Genest St. Isle, France). Food and water were available ad libitum. All procedures were conducted according to the principles and procedures in Guidelines for Care and Use of Laboratory Animals, and validated by local and national ethical committees. Five mice were sacrificed per neurosphere culture. Brains were removed and lateral SVZ dissected under a binocular dissection microscope in DMEM-F12-glutamax 1/50 Glc 45% and incubated with papain, cysteine and DNAse for 30 min at 37°C with pipette dissociation every 10 min to obtain a single-cell suspension. Cells were collected by centrifugation (1000 rcf, 5 min). After resuspension, cells were equally distributed in wells of 96-well plates and cultured in complete culture medium (DMEM-F12 [Gibco], 40 μg/mL insulin [Sigma], 1/200 B-27 supplement [Gibco], 1/100 N-2 supplement [Gibco], 0.3% glucose, 5 mM Hepes, 100 U/ml penicillin/streptomycin) containing 20 ng/mL of EGF and 20 ng/mL of FGF2 (Peprotech), in a 5% CO_2_ environment at 37°C for seven days to obtain primary neurospheres. To test the effect of T_3_ and 4’OH-PPF on neurosphere formation, T_3_ (10nM) and/or 4’OH-PPF (10^−2^mg/L, 10^−1^ mg/L or 3×10^−1^ mg/L) was added to the proliferation medium (12 wells per condition). Medium was renewed every two days.

#### 2.5 Gene expression analysis – mouse neurospheres

After seven days proliferation, neurospheres grown in the same condition were pooled, and RNA was extracted (ThermoFisher RNAqueous MicroKit) and retro-transcriptions performed (Fluidigm) following manufacturers’ instructions. Preamplifications (10 cycles) and qPCR were performed using TaqMan Preamp MasterMix and TaqMan Universal PCR MasterMix, respectively (ThermoFisher), and specific TaqMan probes (References: *Msi1*, Mm00485224_m1; *ActB*, Mm01205647_g1; *Gapdh*, Mm99999915_g1; *Hprt*, Mm00446968_m1) following manufacturer’s instructions. Every qPCR amplification was run in triplicate. ∆Ct have been calculated between target genes and the geometrical mean of three endogenous controls, *ActB*, *Gapdh* and *Hprt*, for each sample. Differences between control and treated groups (ΔΔCt) were used to calculate fold-increases (2^−ΔΔCT^) and to determine changes in target genes expression.

#### 2.6 Chemical exposure - Xenopus laevis tadpoles

The stock solutions of PPF and 4’-OH-PPF were prepared according to following protocol: 15 mg of PPF or 4’-OH-PPF were dissolved in 5 ml of DMSO to create a 3g/L stock solution (stored at −20°C). Final exposure solution was prepared every day of exposure using fresh aliquots of stock solutions (prepared by cascade dilutions from 3g/L stock) by adding 1μL of stock solution to 10 mL of Evian water. All groups including control contained 0.01% DMSO. Fifteen NF 45 TH/bZip eGFP *Xenopus laevis* tadpoles were placed into each well of 6-well plate. Each well corresponds to one exposure group. Eight mL of previously prepared final exposure solution was added into the corresponding well. Tadpoles were exposed for 72 hours at 23°C in dark with daily renewal at the same hour for the XETA assay. For brain gene analysis exposure was limited to 24 hours. Tadpoles were exposed to both PPF and 4’-OH-PPF in the presence and the absence of 5nM T_3_, the most biologically active form of TH.

#### 2.7 XETA - Imaging

At the end of the exposure, tadpoles were rinsed in Evian and anaesthetized using MS-222 (100 mg/L; Stock 1g/L-1g of MS-222 and 1g of sodium bicarbonate dissolved in 1 L of animal water, pH adjusted to pH 7.4-8; stock diluted 10× in animal water). One anaesthetized Tg(*thibz*:GFP) tadpole was placed per well into 96-well plate (black, conic based) and positioned using a fine transfer pipette so the ventral region of the tadpole was facing upwards. To acquire color images we used Olympus AX-70 binocular equipped with 25× objective, long pass GFP filters and a Q-Imaging Exi Aqa video camera. Images were captured using the program QC Capture pro (QImaging) with 3s exposure time. After the image acquisition, tadpoles were euthanized in MS-222 1g/L, fixed overnight in paraformaldehyde 4% and stored at −20°C in cryoprotectant (150g of sucrose and 150 mL of ethylen glycol qsp to 500 mL by PBS 1×). Using the program ImageJ non-specific signal was excluded by splitting the image into 3 layers (red, blue and green channel) followed by subtraction of the red and blue channel from the green one. Integrated density of the green channel was then quantified. Statistical analysis was performed in the program GraphPad Prism 7. Tests used to each experiment are described in the figure legends.

#### 2.8 RT-qPCR – Xenopus laevis tadpole brains

At the end of the 24h exposure, tadpoles were rinsed and anaesthetized in 100 mg/L MS-222. RNA-free dissecting tools were used to dissect tadpole brains. Two brains were placed in one microdissection tube (nuclease free) containing 100 μL lysis solutions from an RNA extraction kit (Ambion RNAqueous). Liquid nitrogen was used to snap freeze the tubes which were then stored at −80°C prior to RNA extraction according to the manufacturer’s instructions. The RIN of each sample was verified using Agilent biaonalyzer. Only samples with RIN higher than 7.5 were used for further RT-qPCR. cDNA was synthetized using Reverse Transcription Master Mix (Fluidigm). The synthetized cDNA was 20× diluted (5μL of DNA in 95μL of nuclease free water). qPCR mix contained following components: 0.15 μL of reverse primer (10pM), 0.15 μL of forward primer (10pM), 1.7 μL of nuclease free water and 3 μL of Power SYBR master mix per reaction. For the final reaction 5 μL of previously prepared qPCR mix were added in one well of 382 well-plate together with 1μL of diluted cDNA. Water control and RT- control was included on every plate. Comparative C_t_ measurements including melt curve were quantified using QuantStudio 6 flex qPCR device (Life technologies). C_t_ value of each sample was exported in excel format. Geometric mean of two housekeeping genes (*ef1a* and *odc*) of each sample was quantified and excluded from the Ct values corresponding to the sample (resulting in ΔC_t_ value). Median values of control group of each gene was calculated and extracted from each ΔC_t_ value to normalize. Fold change using formula =(power 2; - ΔC_t_ value of a sample) was calculated and statistical analysis using resulting values was performed using the program Graphpad Prism7.

#### 2.9 Immunohistochemistry in Xenopus laevis tadpole brains

After 24h of exposure, tadpoles were euthanized using MS-222 (1g/L) and fixed in 4% paraformaldehyde for 3h at room temperature. After the fixing period the brains of the tadpoles were dissected and placed in PBT (1% Triton X-100 in PBS) for at least 18h at 4°C. Next the brains were blocked in 10% (vol/vol) normal goat serum (NGS) in PBT for 3h at 4°C. The primary antibody (Rabbit polyclonal to Msi1-Abcam, UK) was diluted 1/300 in 10% NGS in PBT and the brains were incubated in the solution over night at 4°C. Next morning, the plates were removed to RT for 1h prior to multiple washes in PBT during a period of at least 8 hours. Secondary antibody (Alexa Fluor 488-conjugated anti-rabbit (Thermofisher Scientific) diluted 1/400 in 10% NGS in PBT was applied over night at 4°C. After the secondary antibody treatment, brains were washed several times in PBS during the period of 4 hours. Images of the dorsal side of the brains were taken using Leica MZ16F stereomicroscope equipped by QImaging Retiga-SRV camera. Images were analyzed using the program ImageJ. Region of interest was manually selected and the mean integrated density of each image was calculated with the threshold set to 60.

#### 2.10 Mobility of Xenopus laevis tadpoles

After 72h exposure tadpoles were rinsed in Evian water and placed in 12-well plates with one tadpole per well each containing 4 mL Evian water. Each group was given 10 minutes acclimatization time prior to the trial. Tadpole movements were traced for 10 minutes with 30s light/30s dark intervals using a Daniovision (Noldus) device. Total distance traveled during each 10s of the 10min trials were quantified and exported using the Ethiovision XT 1.5 program. All values were normalized to the mean of the control group of the 0s-10s time period.

#### 2.11 Structural measurements of Xenopus laevis tadpole brains

Tadpoles previously fixed in step 2.7 were placed in a Petri dish and manually positioned to expose the dorsal part of the body. Color images of tadpole heads on black background were taken using a Leica MZ16F stereomicroscope equipped by QImaging Retiga-SRV camera. Brain structures were measured in program ImageJ. MS Excel was used to calculate head/brain ratio.

#### 2.12 Statistical analysis

Statistical analysis was performed using program Graph Pad Prism 7. The D’Agostino & Pearson normality test was done to determine if values follow Gaussian distribution (parametric) or not (non-parametric). One way ANOVA was done in the case of parametric distribution and followed by Dunnett’s multiple comparison post-test. For results displaying non-parametric distribution, Kruskal-Wallis test with Dunn’s multiple comparison post-test was used. If only two groups were compared, t-test (parametric) or Mann-Whitney’s test (non-parametric) was used.

## 3. Results

### 3.1. Comparative binding scores predict a strong steric fit into the T3 binding pocket

A comparative ligand-protein interaction of common insecticides, TH and TH analogues against the human thyroid hormone receptors TRα was carried out and ranked according to their degree of interaction (Supplementary table 1). T_3_, 4’-OH-PPF and PPF ligands show comparative binding scores −62.83 and −76.47 (Fig 1A) and −73.3 (Fig S1) respectively for TRα. These scores predict a strong steric fit of 4’-OH-PPF and PPF, in a similar fashion to T_3_, into the T_3_ binding pocket. With the same method, NH3 displayed comparative binding scores of −78.7 for TRα.

**Figure 1.**
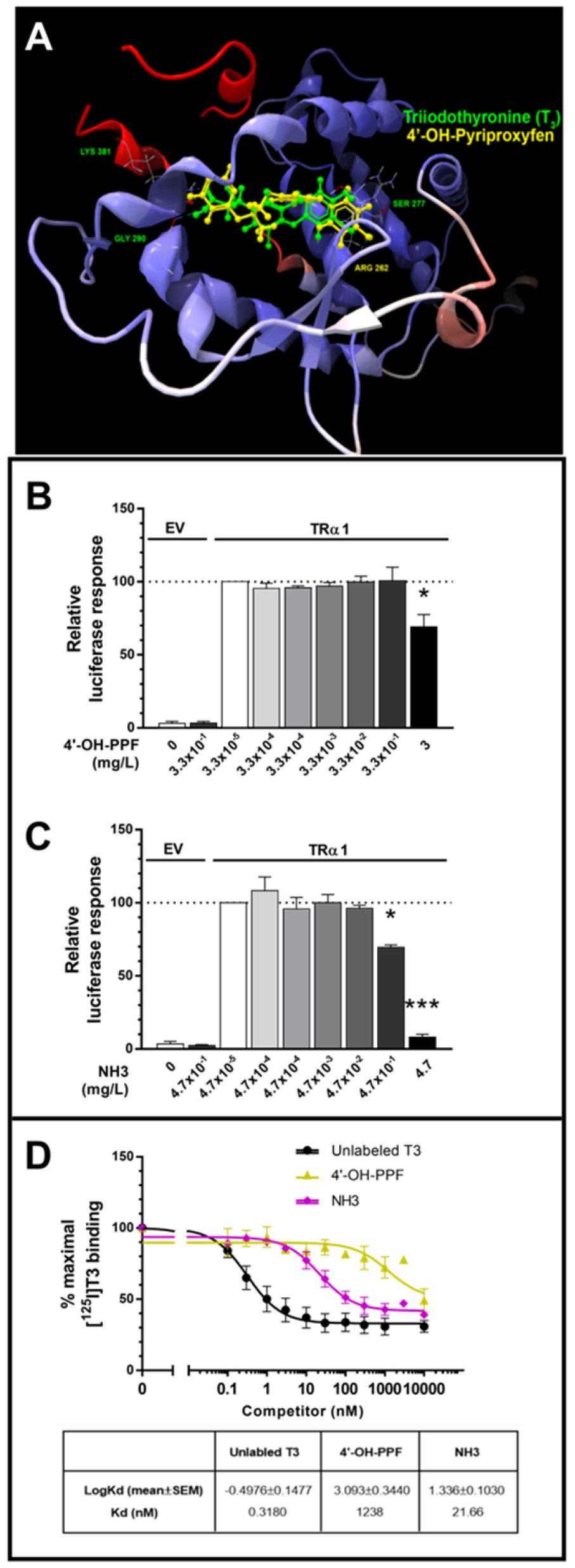
4’-OH-PPF acts as Thrα antagonist with affinity 50× lower than NH3. (**A**) Docking result of Triiodothyronine (T3) and 4’-OH-Pyriproxifen on the crystal structure of human thyroid hormone receptor alpha (TRα, 4LNW). <10,000 iterations were run for each ligand within the T_3_ binding pocket, with top 10 docking results analysed for varying conformations using CLC Drug Discovery 3. Ligand docking of triiodothyronine (T_3_) with three hydrogen bonds are; LYS^381^, GLY^290^ and SER^277^ for TRα. Ligand docking of 4’OH-Pyriproxyfen with one hydrogen bond is ARG^262^ for TRα. T_3_ and 4’-OH-Pyriproxifen ligands show comparative binding scores −65.25 and −77.33 respectively for TRα. These scores predict a strong steric fit of 4’-OH-Pyriproxifen into the T_3_ binding pocket. (**B-C**) The transcriptional activity of TRα1 stimulated by 1 nM T3 was significantly reduced by increasing concentrations of 4’-OH-PPF (**B**) and NH3 (**C**) (EV: empty vector control). The data are presented as mean±SEM of two independent experiments performed in triplicate (One-way ANOVA p<0.05; Tukey’s post-test compared to 0 nM competitor, *p<0.05, **p<0.01, ***p<0.001). (**D**) [^125^I]T3 dissociation curves showing the affinity of TRα1 for 4’-OH-PPF and NH3. High concentrations of 4’-OH-PPF and NH3 reduced TR-bound [^125^I]T3, indicating competitive binding to TRα1. 10000 nM of 4’-OH-PPF and NH3 corresponds to 3 mg/L and 4.7 mg/L respectively. The affinity of TRα1 for 4’-OH-PPF was lower than for NH3, indicated by the right-shift of the curve and higher Kd for 4’-OH-PPF. The data are presented as mean±SEM of at least two independent experiments performed in duplicate.

### 3.2. High concentrations of 4’OH-PPF inhibit T_3_-induced transcriptional activity of TRα1

The effect of 4’-OH-PPF on the T_3_-induced transcriptional activity of TRα1 was determined by an *in vitro* luciferase reporter assay using human placenta JEG3-cell cultures. Transcriptional activity was significantly decreased by the highest concentration of 4’-OH-PPF (3 mg/L) (Fig.1B). NH3 also reduced the transcriptional activity of TRα1, but already at lower concentrations (4.7 and 4.7×10^−1^ mg/L) (Fig.1C). These findings shown 4’-OH-PPF can interfere with T_3_-induced transcriptional activity of TRα1, but only at a high dose, consistent with the low affinity binding to the receptor.

### 3.3. 4’-OH-PPF binds to TRα1 with low affinity

[^125^I]T3 competitive binding assays were performed to determine whether 4’-OH-PPF binds to the TRα1 receptor. Unlabelled T_3_ and NH3, were included as positive controls. The Kd of the NH3 was 60-fold higher than the K_d_ of unlabelled T_3_ (21.6 vs. 0.32 nM), suggesting that NH3 can bind to the TRα1 receptor with a lower affinity than T_3_. Competitive binding of 4’-OH-PPF to TRα1 only occurred at the highest concentration (3 mg/L) and the estimated K_d_ of 4’-OH-PPF (~1.2 μM) was more than 3000-fold higher than T_3_ and 50-fold higher than NH3 (Fig 1D).

### 3.4. PPF and 4’-OH-PPF affect TH signaling in the XETA assay

Transgenic Tg(*thibz*:GFP) tadpoles were exposed to environmentally-relevant dose ranges of 4’-OH-PPF or PPF in the presence or absence of T_3_ (5nM). The concentrations selected varied from 0.1 ng/L through 100ng/l (concentration of PPF detected in the surface water in Spain (Belenguer et al., 2014), 0.01mg/L – PPF drinking water limit according to WHO, up to the WHO acceptable PPF daily intake level (3×10^−1^ mg/L) (WHO, 2007).

PPF, in the absence of T_3_, significantly reduced GFP fluorescence emitted from the tadpole head region at the following concentrations: 10^−5^, 10^−4^, 10^−2^ and 3×10^−1^ mg/L (Fig. S2A). In the presence of T_3_, PPF did not exhibit any thyroid hormone disrupting activity (Fig S2B). In contrast, 4’-OH-PPF was not active without T_3_ (Fig 2B), but in its presence, all concentrations of 4’-OH-PPF significantly reduced the GFP signal (Fig 2C). These results suggest that both PPF and its metabolite 4’-OH-PPF exhibit TH-disrupting properties, but that the parent compound is active in the absence of T_3_ whereas the metabolite antagonizes the effects of T_3_.

**Figure 2.**
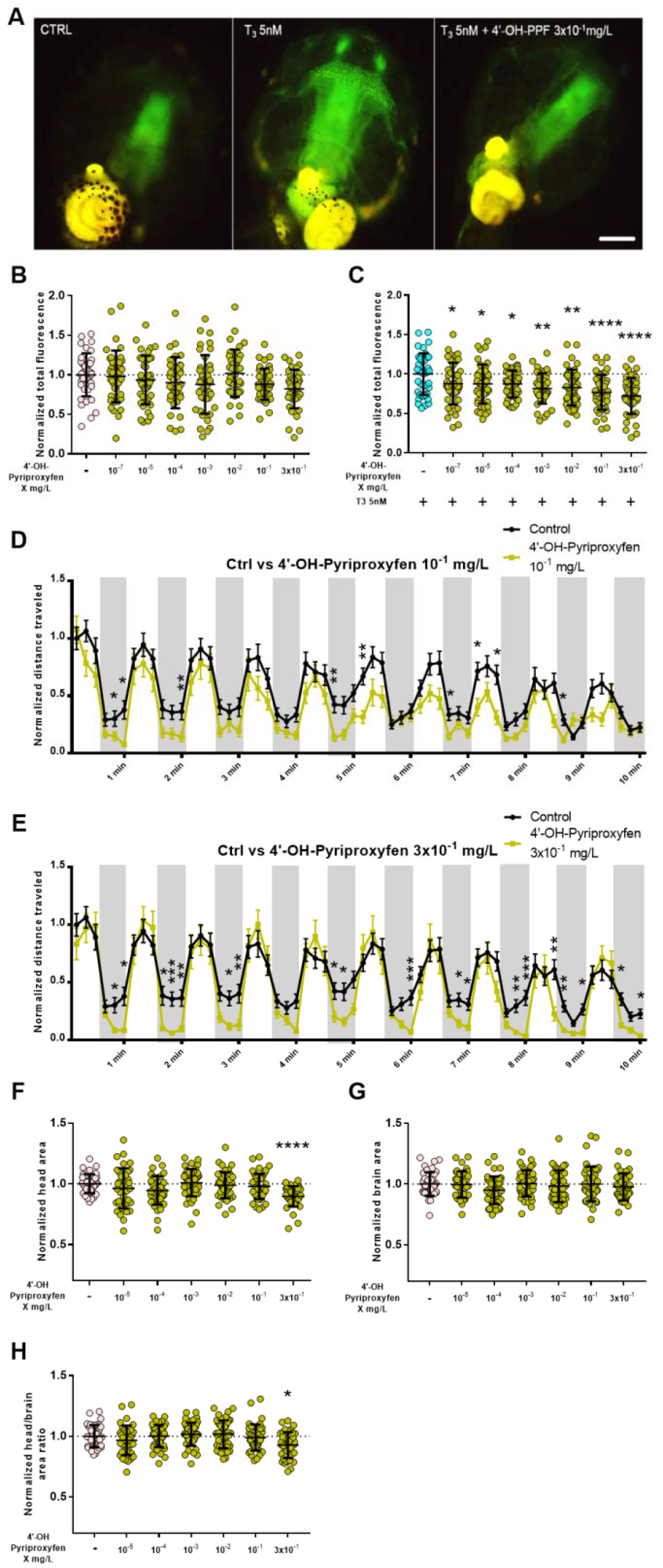
4’-OH-PPF affects thyroid hormon signalling and mobility of *Xenopus laevis* tadpoles. **(A)**Tg(*thibz*:GFP) tadpoles exposued for 72h to (from left to right) vehicle control, T_3_ 5nM and T_3_ 5nM+ 4‘-OH-PPF 3×10^−1^mg/L. Scale bar 500 μm. **(B-C)**72 hours exposure of st 45 Tg(*thibz*:GFP) *Xenopus laevis* tadpoles to 4’-OH-PPF in the absence (**B**) or presence of T_3_ 5nM (**C**). Values normalized to control group (**B**) or T_3_ group (**C**). Pool of 3 independent experiments; n=15 per experiment. One-way ANOVA with Dunn’s post test (Mean ± SDs, *P < 0.05, **P < 0.01, ***P < 0.001, ****P < 0.001). (**D-E**) Total distance travelled by tadpoles exposed to 4’-OH-PPF. 10 minute trial, distance traveled counted every 10 seconds. Dark background represents dark period, white background represents light period. Values normalized to first 10 second period of control group. Pool of 3 independent experiments; n= 12 per experiment. Kruskal-Wallis (Mean ± SEM, *P<0.05, **P < 0.01, ***P < 0.001). (**F-H**) Area of the head (**F**), brain (**G**) and head/brain ration (**H**) of st45 *Xenopus laevis* tadpole exposed to 4’-OH-PPF during 72h. Values normalized to control group. Pool of 3 independent experiments; n=15 per experiment. One-way ANOVA with Dunn’s post test (Mean ± SDs, *P<0.05, ****P < 0.001).

### 3.5. PPF and 4’-OH-PPF affect tadpole mobility

Tadpoles exposed to PPF for 72 h reacted to the light stimulus but moved significantly less than controls (Fig S2C-H). Significant differences were observed in tadpoles exposed to all concentrations of PPF with the exception of 10^−3^ mg/L. In the case of 4’-OH-PPF, the tadpoles exhibited reduced mobility compared to controls, reaching significance at the two highest concentrations (10^−1^ and 3× 10^−1^ mg/L)(Fig 2D,E).

### 3.6. PPF and 4’-OH-PPF affect brain morphology

The total head and brain area was measured using the ImageJ program. Head /brain ratio was calculated using MS Excel. PPF did not affect brain or head size, nor was the head/brain size ratio significantly different compared to controls (Fig S4C-E). However, tadpoles exposed to 3×10^−1^ mg^/^L of 4’-OH-PPF exhibited significantly smaller heads compared to control (Fig 2F), thus resulting in biased brain/head ratios (Fig 2H), however the brain size of these tadpoles was not affected (Fig 2G). The width and length of brain compartements were meassured (Fig S4B). The width of forebrain (H1) was increased in tadpoles exposed to PPF at concentrations 10^−4^ mg/L – 10^−1^ mg/L (Fig S4F). The width of midbrain (H3) was significantly increased in tadpoles exposed to concentrations of PPF 10^−3^ mg/L and higher (Fig S4H). The length of the whole brain (L3) was significantly increased by exposure to PPF at concentration 10^−3^ mg/L and higher (Fig Fig S4H). In the case of 4’-OH-PPF, the lowest concentration 10^−5^ mg/L significantly increased the width of forebrain and midbrain junction (Fig S4M) and the width of midbrain (Fig S4N).

### 3.7. PPF and 4’-OH-PPF affect brain musashi-1 gene and protein expression in Xenopus laevis tadpoles

We studied the consequences of a 24h exposure with PPF or 4’-OH PPF on brain gene *musashi-1* (*msi1*) expression and its encoded protein in Xenopus *in vivo*. *Msi1* brain gene and protein expression was analyzed following 24h exposure to each compound as a function of results seen in the XETA test. Thus, PPF was run without T_3_ whereas 4’-OH-PPF was tested in the presence 5nM T_3_. T_3_ 5nM significantly reduced the expression of *msi1* gene and its encoded protein (Fig 3 A-C). 4’-OH-PPF increased the expression of *msi1* gene at both concentrations tested in the presence of T_3_ 5nM (Fig 3B) and increased msi1 protein expression at 10^−2^ mg/L (Fig 3C). PPF did not affect *msi1* gene expression (Fig S3A) however the highest concentration 3×10^−1^ mg/L significantly reduced the expression of msi1 protein in the brain of NF45 tadpoles after 24 hours (Fig S3B).

**Figure 3.**
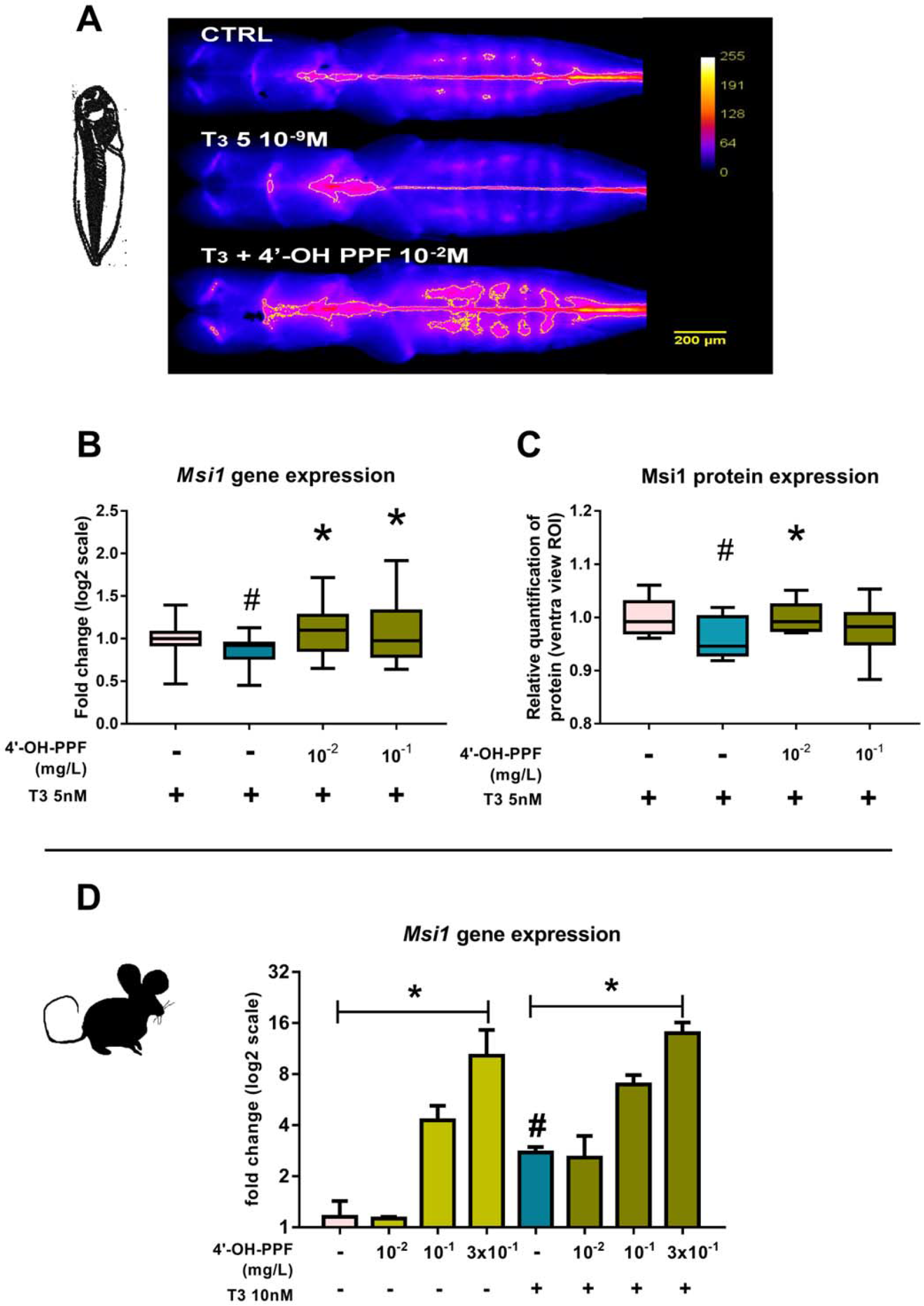
*Musashi1* gene expression and expression of its encoded protein is modified by TH and 4-OH-PPF. (**A**) Representative pictures of immunohistochemistry realized on solvent control, T_3_ 5nM and T_3_+ 4’-OH-PPF 10^−2^ mg/L, ROI is delimited by thin white line. (**B**) *Msi1* gene expression on dissected brains from NF46 tadpoles exposed for 24h to 4’-OH-PPF (10^−2^; 10^−^1 mg/L) in challenge with T_3_ 5nM. (**C**) Msi1 protein expression on dissected brains from NF46 tadpoles exposed for 24h to 4’-OH-PPF (10^−2^; 10^−1^ mg/L) in challenge with T_3_ 5nM. Antibody Alexa Fluor 488-conjugated anti-rabbit. Quantification of the protein level based on fluorescence intensity from dorsal views. Two independent experiments in case of protein expression study with 4 to 10 tadpoles per experiment were done. Three independent experiments in the case of gene expression study, n=5 per experiment. (**D**) *Msi1* gene expression in mouse neurospheres after 7 days of proliferation. Graphs represent the pool of the experiments (each normalized to its own control). Non-parametric ANOVA; Kruskal-Wallis with Dunnet’s post test. *p<0.05. Statistics comparing T_3_ with control - Mann-Whitney; # p<0.05.

### 3.8. 4’-OH-PPF increases Msi1 gene expression in mouse neurospheres

*Msi1* gene expression in mouse neurospheres after 7 d proliferation in the presence of 4’-OH-PPF was analyzed by RT-qPCR. The expression of *Msi1* gene was increased 3-fold by T_3_ 10 nM and the highest concentration of 4’-OH-PPF 3×10^−1^ mg/L in the presence and absence of T_3_ 10 nM (fold induction 10 and 14 respectively) (Fig 3D).

## 4. Discussion

Given paucity and the contradictory nature of studies on the potentially adverse effects of PPF, we investigated whether its main metabolite, 4’-OH-PPF, affects TH signaling and *Msi1* gene expression using *in silico*, *in vitro* and *in vivo* approaches. TH modulates every stage of brain development, thus any modification of TH availability and action could have direct consequences on neurogenesis and brain morphology, as well as on cranio-facial development (Bernal, 2005). The importance of TH for brain development, and notably for regulating *Musashi 1* gene expression in neuroprogenitor cells (López-Juárez et al., 2012), raised the question as to whether TH disruption caused by PPF exposure might be implicated in the severity of the incidence of microcephaly in certain regions of Brazil, especially North Eastern Brazil (area with the highest incidence of microcephaly), where it has been used intensively (Parens et al., 2017).

The *in silico* results revealed that 4’-OH-PPF had a high steric fit for both TRα we tested whether the metabolite blocked T_3_ binding to the TR LBDs. The *in vitro* study showed that 4’-OH-PPF acts as TRα antagonist at a concentration 3 mg/L, approximately 10× the WHO recommended daily intake of PPF (WHO, 2007) (Fig. 1B,D). These findings indicate that although TRα1 can bind 4’-OH-PPF, the affinity is much lower than for T_3_ and NH3, which is not in accordance with predicted strong steric fit of 4’-OH-PPF into the T_3_ binding pocket as shown by *in silico* docking.

However, our findings that 4’-OH-PPF affects expression *msi1* gene and protein offer a plausible mechanism to explain potential increase of susceptibility to ZIKV. We show that 4’-OH-PPF increased the levels of *msi1* gene and its encoded protein in the brain area of early stage tadpoles in the presence of T_3_ 5nM and increased the expression of *Msi1* gene in adult mouse neurospheres in the presence and absence of T_3_ 5nM. *Msi1* RNA has been shown to interact with the ZIKA RNA by mediation of the 3’UTR of the ZIKA virus and is required for ZIKA replication (Chavali et al., 2017). Msi1 is an important translational regulator in neuronal stem cells in both vertebrates and invertebrates which is enriched in neural progenitors at the ventricular and sub ventricular zones but absent from the cortical plate (Chavali et al., 2017). We had previously shown that this gene is downregulated *in vivo* in mice subventricular zone by T_3_/TRα during progenitor to neuroblast commitment (López-Juárez et al., 2012). We analyzed the regulatory element upstream of human *MSI1* gene by NHR scan, finding that this sequence contains a number of DR4 type response elements, i.e. potential TR binding sites (Chang and Pan, 1998; Paquette et al., 2014; Quack et al., 2002). A recent study compared neuroprogenitor cells (NPCs) from twins discordant for Congenital Zika Syndrome (CZS) displayed differences in NPC gene expression that could underly increased susceptibility to ZIKV-induced microcephaly through viral replication (Caires-Júnior et al., 2018).

Arguments against the implication of PPF in microcephaly have been put forward by the manufacturer, Sumitomo. In their most relevant study, Sumitomo exposed pregnant rat dams during the days 7 to 17 of gestation to dose range of PPF. The pups were then observed for physical deformations and their organs weighed. According to the producer, results did not indicate any significant changes on the nervous or reproductive systems (Saegusa, 1988). However a recent review (Parens et al., 2017) suggests that the assay may not be sufficiently sensitive and was largely restricted to analyzing impact on adult animals. Parens et al, also noticed that the report describes low brain mass and one case of microcephaly. This case was considered irrelevant by Sumitomo who argued that microcephaly should not be considered due to its dose-dependence. Parens et al., also proposed a possible mechanism of action for PPF-driven microcephaly implicating cross reactivity of PPF with retinoic acid (RA) signaling. Juvenile hormones and retinoic acid share structural similarity and it is known, that some juvenile hormones are able to bind to the retinoid acid receptor (RAR) (Harmon et al., 1995; Palli et al., 1991). They propose that PPF, a juvenile hormone analog, binds to RAR and mimics the effect of RA, which itself has been previously connected with the induction of microcephaly (Soprano and Soprano, 1995). TH and RA interact during neurogenesis (Gil-Ibáñez et al., 2014), with RA and TR binding to nuclear receptors of steroid/thyroid hormone receptor superfamily, sharing certain gene response elements (Glass et al., 1989) and their heterodimeric retinoid X receptor (RXR) partner on DNA (Kliewer et al., 1992; Zhang and Kahl, 1993). It has been also demonstrated that RA and TH interact to regulate craniofacial development with RXR partially mediating their interaction (Bohnsack and Kahana, 2013).

Only two studies, both using Zebrafish, have addressed the question of whether PPF could be implicated in microcephaly, each producing contradictory results (Dzieciolowska et al., 2017; Truong et al., 2016). One study (Truong et al., 2016) demonstrated that PPF induces adverse morphological changes including craniofacial defects in zebrafish at 5.2 μM (1.66 mg/L) - EC50. In the behavioral test PPF at 6.4μM (2 mg/L) and 64μM (20mg/L) significantly reduced the mobility of the zebrafish larvae compared to control animals. The authors concluded that the developmental toxicity of PPF may not be limited only to insects (Truong et al., 2016).

## Conclusions

Taken together our results and previous findings we provide a range of evidence, that PPF and its metabolite 4’-OH- PPF affect TH signaling, brain gene expression and behavior. We propose a plausible mechanism of action, ie that changes in *musashi1* expression could exacerbate ZIKA virus infection.

## Acknowledgements

We are grateful for the work in the Xenopus facility of Gérard Benisti, Philippe Durand and Jean-Paul Chaumeil. We would like to thank all our colleagues for their comments on the manuscript, and Fiene Kuijper for analysis of the mouse neurosphere data. This work was supported by grants from Centre National de la Recherche Scientifique (CNRS), Muséum National d’Histoire Naturelle (MNHN), PNREST THYPEST EST-2014-122 and European Union contract EDC MIX RISK_GA N°634880.

## Author Contributions

P.S., J.B.F and B.D. designed the study. P.S. carried out the *in vivo* experiments and analyzed the data. S.L. helped with the immunohistochemistry. K.W. and T.V. carried out the *in vitro* experiments. B.M. carried out the *in silico* study. M.L. helped with the fluorescence read-out, immunohistochemistry, brain dissections and prepared Xenopus breedings. P.S, J.B.F., and B.D. wrote the paper. All authors discussed the results and commented on the manuscript.

